# Efficient small fragment sequencing of human, cow, and bison miRNA, small RNA or csRNA-seq libraries using AVITI

**DOI:** 10.1101/2024.05.28.596343

**Authors:** Anna L. McDonald, Andrew M. Boddicker, Marina I. Savenkova, Ian M. Brabb, Xiaodong Qi, Daniela D. Moré, Cristina W. Cunha, Junhua Zhao, Sascha H. Duttke

## Abstract

Next-Generation Sequencing (NGS) catalyzed breakthroughs across various scientific domains. Illumina’s sequencing by synthesis method has long been essential for NGS but emerging technologies like Element Biosciences’ sequencing by avidity (AVITI) represent a novel approach. It has been reported that AVITI offers improved signal-to-noise ratios and cost reductions. However, the method relies on rolling circle amplification which can be impacted by polymer size, raising questions about its efficacy sequencing small RNAs (sRNA) molecules including microRNAs (miRNAs), piwi-interacting RNAs (piRNAs), and others that are crucial regulators of gene expression and involved in various biological processes. In addition, capturing capped small RNAs (csRNA-seq) has emerged as a powerful method to map active or “nascent” RNA polymerase II transcription initiation in tissues and clinical samples. Here, we report a new protocol for seamlessly sequencing short DNA fragments on the AVITI and demonstrate that AVITI and Illumina sequencing technologies equivalently capture human, cattle (*Bos taurus*) and the bison (*Bison bison*) sRNA or csRNA sequencing libraries, augmenting the confidence in both approaches. Additionally, analysis of generated nascent transcription start sites (TSSs) data for cattle and bison revealed inaccuracies in their current genome annotations and highlighted the possibility and need to translate small RNA sequencing methodologies to livestock. Our accelerated and optimized protocol therefore bridges the advantages of AVITI sequencing and critical methods that rely on sequencing short DNA fragments.

## INTRODUCTION

Next-generation sequencing (NGS) revolutionized biology and biomedicine, and has led to considerable advancements in research, clinical diagnostics, agricultural and environmental applications. Recent key contributing factors included cost-efficient sequencing, greater accessibility for researchers and clinicians, increased speed, throughput, and precision. Simultaneous analysis of numerous sequences facilitated the identification of genetic variants, aiding understanding of diseases, population genetics, breeding, and evolutionary studies.

The Illumina sequencing by synthesis method has long been a cornerstone in NGS, but new technologies are now emerging. Recently, Element Biosciences released their AVITI platform. Instead of sequencing by fluorescently-labeled and reversibly-terminated nucleotides, as done by Illumina, AVITI circularizes sequences and uses specific detector molecules called avidites. As these multivalent molecules are highly specific and bind multiple extension sites within an amplified “polony”, AVITI can use less reagents which translates into low sequencing costs and less background signal (Arslan et al. 2024). However, DNA circularization may be size-dependent and generally inefficient for shorter polymers, depending on circularization mechanism, especially below 150 bp (Jacobson and Stockmayer 1950; Joffroy et al. 2018; Gouzouasis et al. 2023).

Small RNAs (sRNAs) such as micro RNAs (miRNAs), small interfering RNAs (siRNAs), piwi-interacting RNAs (piRNAs), and others, are crucial regulators of gene expression, involved in various biological processes including development, defense against viruses and transposons, and the maintenance of genome stability (Chen 2009; Law and Jacobsen 2010; Gouzouasis et al. 2023). Consequently, they are a fundamental area of study in molecular biology and a focus in the search for future therapeutic interventions. In addition, capturing capped small RNAs (csRNA-seq) has emerged as a powerful method to identify sites of active or “nascent” transcription from total RNA or clinical samples (Duttke et al. 2019; Lam et al. 2023) and circulating cell-free DNA (cfDNA), which is typically around 180-200 bp in length, are of emerging interest for diagnostics, disease monitoring, and therapeutic applications (Gao et al. 2022). We therefore compared the AVITI and Illumina sequencing technologies in their ability to capture sRNA sequencing libraries.

Here we present two expedited and refined protocols for short fragment sequencing with AVITI, which align seamlessly with the majority of commercially available sRNA library kits. We show that sequencing short fragments like small RNAs (18-60 nt in size) or initiating RNA polymerase II transcripts (csRNA-seq) gives uniform results with AVITI and Illumina sequencing technologies. Moreover, generation of sRNA and the first csRNA-seq libraries in cattle and bison demonstrate the applicability of our approach in livestock. Our analyses reveal that 5’ annotations of many Reference Sequence annotations (RefSeq) of cattle and bison, but not humans, are often inaccurate, highlighting the importance of the provided experimental data as, among others, accurate TSSs are critical for successful genome engineering approaches (Radzisheuskaya et al. 2016; Shamie et al. 2021).

## RESULTS

### New protocols for uniform small RNA coverage among Illumina and AVITI sequencing technologies

To overcome prior limitations, we developed and tested two optimized and accelerated protocols for AVITI short read circularization-based sequencing (Fig. S1) using human cancer cells and blood from cattle and bison, two animals of agricultural importance, and compared the results to Illumina. We generated small RNA (18-60 nt), containing all mono, di, tri, 5’ capped or otherwise 5’ modified sRNAs (Lister et al. 2009)) and csRNA-seq libraries, which are associated with active initiation of RNA polymerase II promoters and enhancers (Duttke et al. 2019; Duttke et al. 2022a). Libraries were then sequenced on the Illumina NovaSeq 6000, Illumina NextSeq 2000, or the Element Biosciences AVITI platforms (Fig. 1), downsampled to an equal number of reads (Table S1), and subsequently compared.

**Fig. 1:**
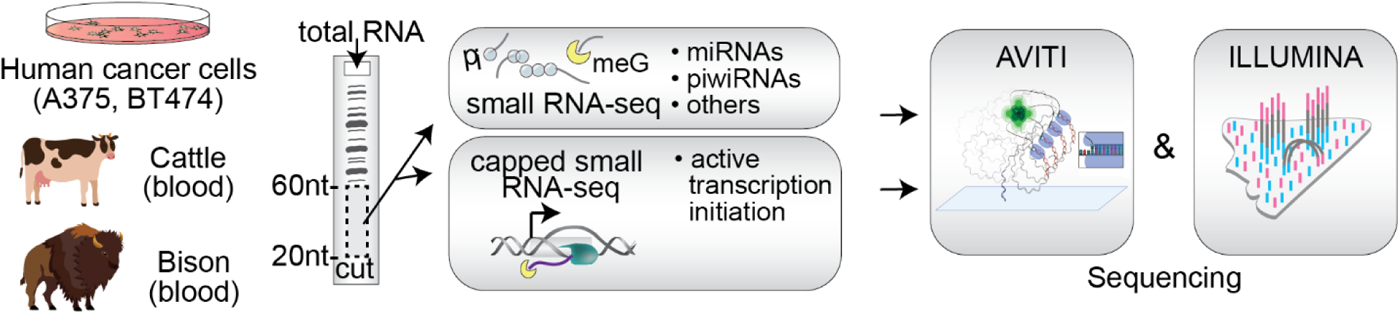
Study design. Small RNAs sizes 18-60 nt were purified from total RNA isolated from human cancer cell lines (A375, BT474), cattle and bison blood. All sRNAs (sRNA-seq) as well as 5’meG cap-enriched RNAs (csRNA-seq) libraries were generated and sequenced on the AVITI and Illumina NGS platforms.

To assess potential biases, we first compared the read length distribution of sequenced fragments in human A375 and BT474 cancer cells in replicate (please see materials and methods). Samples were circularized either on benchtop or directly on the AVITI flow cell prior to sequencing, for the A375 and BT474 cells, respectively. Any technical differences between the Illumina and AVITI methods, such as bridge amplification versus rolling circle amplification, should result in a linear size bias relationship across all samples. By contrast, differences arising from experimental replicates could be non-linear.

Observed differences in size distribution among the two sequencing methods did not correlate with sequenced fragment length for sRNA or csRNA-seq (Fig. 2 A,B, Fig. S2 A,B). Indeed, differences were more pronounced between replicates on each platform than between Illumina and AVITI sequencers (Fig. S3 A,B). Concordantly, expression levels of >5,682 sRNAs as well as >70,351 active regulatory elements, such as promoters and enhancers, captured by csRNA-seq were highly similar across sequencing platforms (r= 0.999 and r= 0.9996, respectively, Fig. 2 C,D, Fig. S3 C,D). Analyzing differentially expressed loci between platforms using DEseq2 (Love et al. 2014) revealed only 10 downregulated protein coding genes in Illumina data, and 28 in AVITI data (log_2_FC, FDR 0.05). Equivalently high correlations were observed among nucleotide frequencies (Fig. S2 C,D), miRNAs (21-24nt in length) and the >55nt long small-nucleolar RNAs (snoRNAs, (Dupuis-Sandoval et al. 2015), Fig. 2 E, Fig. S3 E).

Similarly, no significant differences were observed between sequencing platforms in average GC content of samples, while the average trimmed read length of AVITI reads were slightly lower (Table S1). Average estimated quality across all bases of trimmed reads was higher from AVITI reads (Q=42.4) than Illumina (Q=35.4) (Table S1). Together, these data argue that short fragments are efficiently captured by both sequencing platforms.

**Fig. 2:**
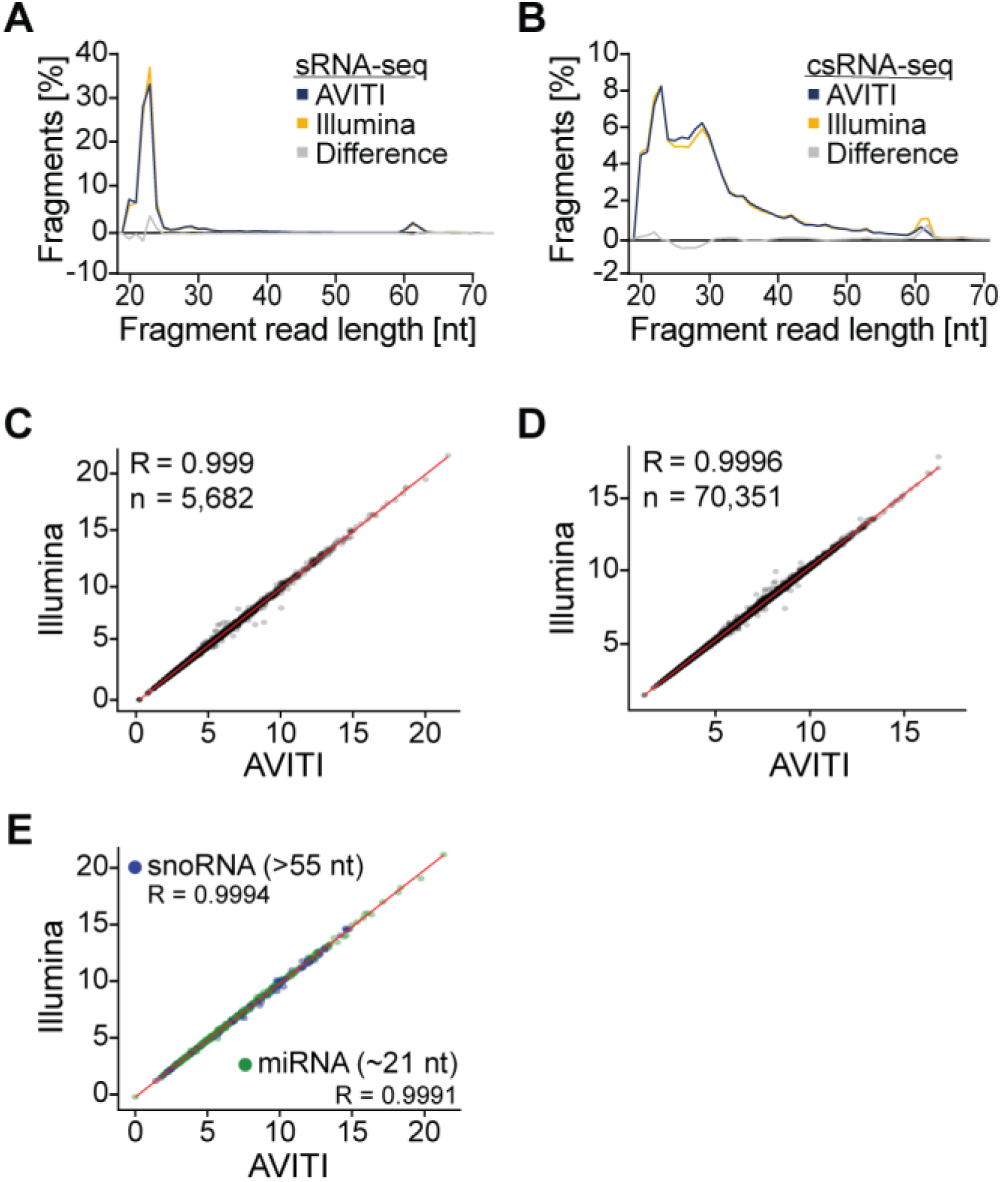
Uniform sequencing of small and capped small RNA-seq libraries on the Illumina and AVITI platforms. **A**. Read length distribution plots of A375 small RNAs sequenced natively on the Illumina and after benchtop circularization (Adapt Rapid PCR-free) on the AVITI platform. The area under each line sums to a total of 100%. Differences between Illumina and AVITI are plotted in grey. **B**. Read length distribution plots of A375 cells capped small RNAs. **C**. Scatterplot comparing the expression level of small RNAs and **D**. capped small RNAs using the Illumina and AVITI platform. **E**. Comparison of the detection of small RNA types of different lengths (miRNAs: 21-24; snoRNAs: 55-61).

### Accurate coverage of small RNAs and active TSSs in cattle and bison reveals a need to improve annotations in livestock

To test our protocol in species of agricultural importance we next performed sRNA and csRNA-seq on blood collected from cattle and bison. For AVITI sequencing, libraries were circularized directly on the flow cell surface. Consistently, we observed similar size profiles and sequencing distributions with either Illumina or AVITI sequencing methods (Fig. S4).

During our analyses we noticed inconsistencies among our experimental transcription start site (TSS) data and many annotated 5’ ends of genes in cattle and bison but importantly, not human (Fig. 3 A-C, Fig. S5 A,B; human: GCF_000001405.40, cattle: GCF_002263795.2, bison: modified_GCA_018282365.1). Most TSSs in cattle and bison were within 100 bp of RefSeq 5’ annotations, 39.7% and 45.5%, respectively, suggesting quality annotation of coding regions. However, genic 5’ ends and promoters were inaccurate. As accurate 5’ annotations are an essential part of many analyses including genome engineering (Radzisheuskaya et al. 2016; Shamie et al. 2021) or decoding gene regulatory programs (Shields et al. 2021), and, to our knowledge, our study presents the first nascent TSSs data for cattle and bison, we investigated the differences further. Computational prediction of annotation 5’ ends and, to some extent, mapping the 5’ ends of steady state rather than nascent transcripts, has been shown to lead to annotation inaccuracies (Affymetrix 2009; Ivanov et al. 2021). It is important to note that TSSs change across tissues and thus no single annotation can accurately capture all genetic variants across all cell types (Shamie et al. 2021). However, annotating unbiased biological features, such as core promoter elements or TSS-proximal nucleotide frequencies, revealed a clear improvement of our experimental TSS over RefSeq. Core promoter elements anchor and dictate the site of RNA polymerase initiation. Consequently, they are highly positionally enriched: the TATA box from ∼-31 to -26, the Initiator from -2 to +4 (Grosveld et al. 1981; Smale and Kadonaga 2003). In addition to the increased information content in the TSS-proximate nucleotide frequencies (Fig. S6), the TATA box and Initiator core promoter elements were found at the expected positions respectively in the human RefSeq and in our experimental TSS data for A375 and BT474, but not in the cattle and bison RefSeq annotation (Fig. 3 D-I, Fig. S5 C,D). Together, these observations provide an independent validation for our experimental TSSs data, stress their importance, and the need to improve genome annotations.

**Fig. 3:**
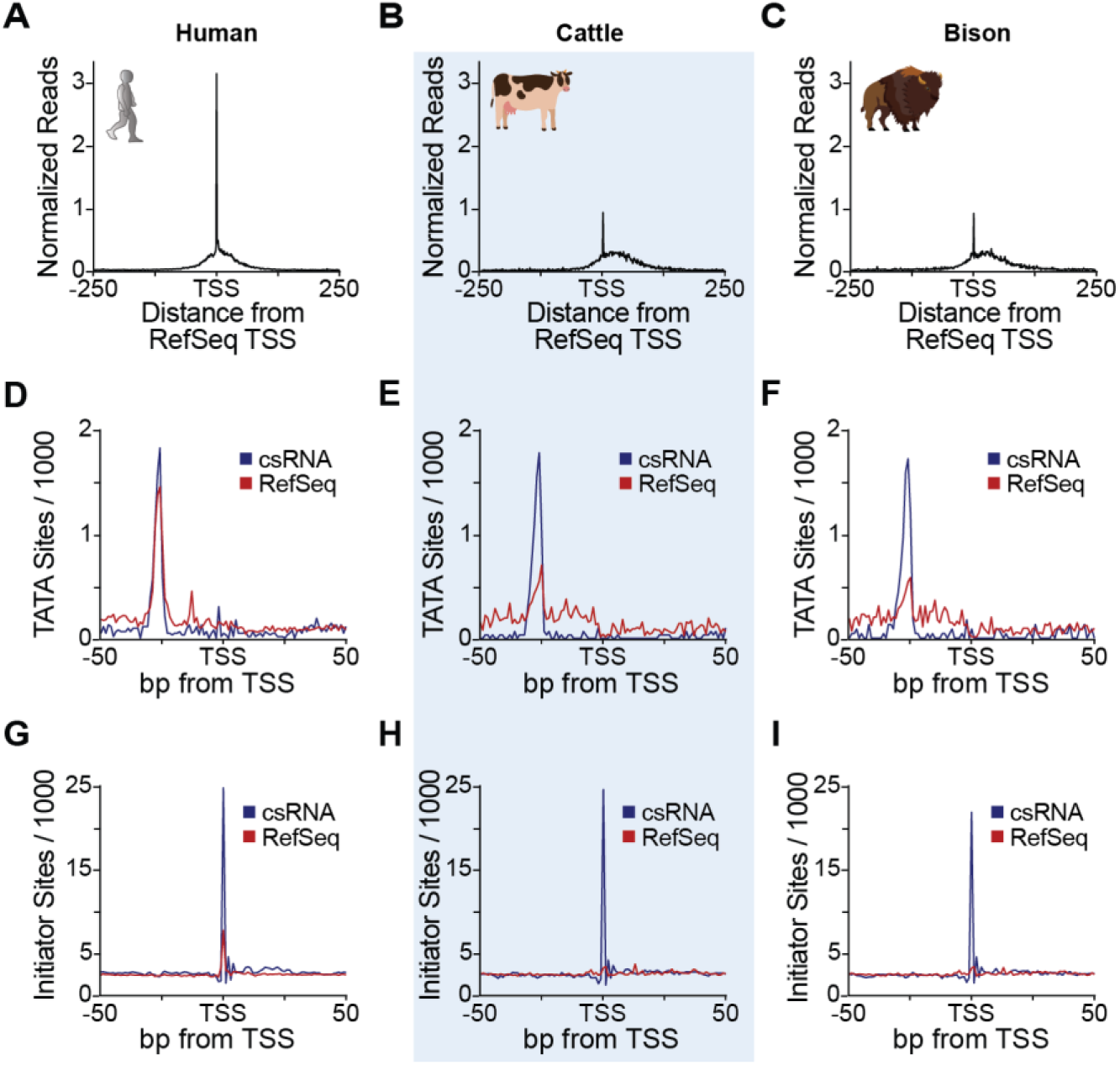
csRNA-seq facilitates improved annotation of livestock genomes. **A.** Comparison of experimentally defined TSSs from human A375 cancer cells, **B.** cattle, and **C.** bison by csRNA-seq relative to the RefSeq annotation. **D.** Comparison of the frequency of TATA box sites per 1000 bp between our experimental TSS and RefSeq for human, **E.** cattle, and **F.** bison. **G.** Comparison of the frequency of Initiator sites per 1000 bp between our experimental TSS and RefSeq for human, **H.** cattle, and **I.** bison.

## DISCUSSION

Here, we provide two optimized and accelerated protocols to circularize and sequence short DNA sequences such as from miRNAs, siRNAs, piwiRNAs, or csRNAs on the AVITI platform and demonstrate uniform results to Illumina sequencing. The required library circularization can be performed on the benchtop before sequencing, or directly on the flow cell surface after loading the linear library (Fig. S1). The benchtop circularization method allows for additional quality control of pre-circularized libraries and includes the addition of a 48 bp backbone sequence, which may mitigate circularization or amplification bias reported of smaller molecules (Jacobson and Stockmayer 1950; Joffroy et al. 2018; Gouzouasis et al. 2023). Circularization directly on the flow cell surface without backbone insertion eliminates any additional hands-on time or quality control steps and shows highly concordant fragment sizes with Illumina data (Fig. S2, Fig. S4). The rolling circle amplification strategy on AVITI is adaptable to a wide range of library molecule sizes ranging from inserts >1000 bp (Carroll et al., *in preparation*) to 20bp, as demonstrated here.

In addition, our study generated sRNA and nascent TSS data for human A375 and BT474 cells, as well as the first data of such kind for cattle and bison, thereby enriching publicly available resources for the scientific community. These data not only demonstrate the utility of csRNA-seq and AVITI sequencing in livestock but also their necessity. While annotated 5’ ends of genes largely agreed with nascent TSSs in well studied organisms, clear differences were observed in cattle and bison. Our experimental TSSs improve upon RefSeq by revealing biological features such as position-constrained core promoter elements (Juven-Gershon and Kadonaga 2010; Haberle and Stark 2018). This is also important as accurate TSSs and targeting of the promoter is critical for genome engineering efforts (Ceasar et al. 2016; Shamie et al. 2021).

The use of multivalent avidites to detect bases on AVITI further resulted in higher sequencing quality metrics compared to Illumina (Table S1). However, sequence polymorphisms among utilized cell lines, or individual bison or cattle, and the specific reference genomes dominated alignment rates, making this difference negligible for small RNAs. Therefore, our study not only provides a new protocol to sequence small nucleotide polymers on the AVITI and nascent TSSs for bison and cattle, but also highlights the possibility to study these small molecules on either platform, increasing the flexibility for researchers, and, by demonstrating uniformity, validating both methods.

## MATERIALS AND METHODS

### Cell culture, siRNA & mRNA transfections

A375 and BT474 cells were grown at 37°C with 5% CO_2_ in DMEM (Cellgro) supplemented with 10% FBS (Gibco), and 50 U Penicillin and 50 μg Streptomycin per ml (Gibco). For total RNA isolation, cells were washed once with ice cold DPBS (Gibco), rested on ice for 5 minutes, washed one more time with ice cold DPBS and then lysed in 1 ml TROZOL. RNA isolated as described by the manufacturer.

Cattle and bison samples were obtained from experimental animals housed at Washington State University, Pullman, WA and University of Wyoming, Laramie WY, respectively. 2.8 ml blood was collected from healthy animals using PAXgene Blood RNA tubes (BD Bioscience) by venipuncture (jugular) and transported on ice to the laboratory for processing. RNA was isolated using the PAXgene Blood miRNA Kit (Qiagen) as described by the manufacturer.

### sRNA and csRNA-seq library generation

Small and capped small (cs)RNA (Duttke et al. 2019) libraries were generated exactly as described in (Duttke et al. 2022b). Small RNAs of ∼20-60 nt were size selected from total RNA by denaturing gel electrophoresis. 10% of these RNAs were decapped and polyphosphates reduced to monophosphates using RppH (NEB) to sequence all small RNAs (sRNA-seq).

The remainder of the size selected sRNAs was enriched for 5’-capped RNAs. Monophosphorylated RNAs were selectively degraded by 1 hour incubation with Terminator 5’-Phosphate-Dependent Exonuclease (Lucigen). Subsequently, RNAs were 5’dephosporylated through 90 minutes incubation in total with thermostable QuickCIP (NEB) in which the samples were briefly heated to 75°C and quickly chilled on ice at the 60 minutes mark. Small RNA and csRNA-seq libraries were prepared using the NEBNext Small RNA Library Prep kit with an additional RppH step (Hetzel et al. 2016), amplified for 13 cycles and sequenced SE80 on the Illumina NextSeq 2000, PE100 on the Illumina NovaSeq 6000 or PE75 on the AVITI.

### AVITI library conversion for sequencing

Libraries prepared from human A375 cells were converted for sequencing on AVITI by following the ‘Rapid Adept PCR-free’ protocol (Element Biosciences, #830-00007, provided also in the supplement of this paper as “Supplemental protocol”).

In brief, two A375 libraries were pooled and 0.15 pmole linear library was denatured and hybridized to a splint oligo mix. Circularization is achieved by ligation of both library ends to a 48bp backbone oligo sequence to form a ssDNA circular molecule. Residual linear library and splint oligo are enzymatically digested, and the reaction is stopped with an EDTA solution. The protocol utilizes a stop solution over a bead-based cleanup to prevent loss of the carefully size selected sRNA and csRNA libraries.

Libraries prepared from human BT474 cells and cattle and bison blood were diluted (8.4-11.2 fmole per flow cell) and loaded directly to the instrument for circularization on the flow cell surface by the AVITI system using the “Cloudbreak Freestyle” chemistry. This process does not incorporate additional sequence to the final library.

All linear libraries were quantified by Qubit dsDNA HS assay (ThermoFisher, Carlsbad,CA) paired with fragment size analysis using Tapestation D5000 High Sensitivity screentapes (Agilent, Santa Clara, CA). Circular libraries were quantified using a qPCR assay as part of the ‘Rapid Adept PCR-free’ protocol (Supplemental protocol, Element Biosciences, #830-00007).

### Sequencing

Illumina NextSeq 2000 was performed at the Washinton State University Molecular Biology and Genomics Core, NovaSeq S6000 sequencing at UC San Diego’s IGM core.

Libraries were sequenced on the AVITI platform at Element Biosciences (San Diego, CA) using 2x76 cycle paired-end sequencing. Libraries prepared from human A375 cells were sequenced using the Cloudbreak chemistry kit (Element Biosciences, #860-00004). Libraries prepared from human BT474 cells, bison, and cattle samples were sequenced using Cloudbreak Freestyle chemistry kits (Element Biosciences, #860-00015) with the modified recipe for short fragments (Supplemental protocol, Element Biosciences, #830-00007).

Custom sequencing primers were added for Read2 and Index1 on the AVITI system to all sequencing runs. Primers were ordered from IDT (Coralville, IA) with HPLC purification (Read2: 5’-GTGACTGGAGTTCCTTGGCACCCGAGAATTCCA-3’, Index1: 5’-

TGGAATTCTCGGGTGCCAAGGAACTCCAGTCAC-3’) and spiked-in to the existing sequencing primer tubes at a final concentration of 1uM following the AVITI user guide (Element Biosciences, #MA-00008).

### Data analysis

Small (s)RNA-seq and capped small (cs)RNA-seq (∼20-60 nt) sequencing reads were trimmed of their adapter sequences using HOMER (homerTools trim -3 AGATCGGAAGAGCACACGTCT -mis 2 -minMatchLength 4 -min 20) (Heinz et al. 2010). To achieve equal read depth, fastq files were subsampled using SeqKit’s sample (version 2.5.1) (Shen et al. 2016) before alignment to the appropriate reference genome: STAR for human (Dobin et al. 2013) and Hisat2 for livestock (default parameters) (Kim et al. 2019). Alignment files (.sam) were converted into tag directories using HOMER2 (batchMakeTagDirectory.pl sam_infofile.txt -cpu 8 -genome {species} -omitSN - checkGC -single -r). Features (peaks), representing strand-specific loci with significant transcription initiation (Transcription Start Regions, TSRs) for csRNA-seq or expressed small RNAs for sRNA-seq, were defined using HOMER’s findcsRNATSR.pl and findPeaks, respectively. A minimum read count of 10 per 10 million was required for regions to be considered (findcsRNATSS.pl csRNA -o output_dir -i sRNA -genome [species] -ntagThreshold 20 -cpu 30, findPeaks {csRNA} -o output_dir -i csRNA -gtf [species] -style tss -ntagThreshold 20). Small RNA-seq data were integrated into the csRNA-seq analysis to eliminate loci with csRNA-seq signal arising from non-initiating, high abundance RNAs captured by the method. Replicate experiments were pooled into meta-experiments for each condition before identifying features. Additional information and analysis tutorials are available at https://homer.ucsd.edu/homer/ngs/csRNAseq/index.html.

### Differential Expression

The previous TSRs that were combined across replicates were used to create new merged files for pairwise comparisons using HOMER’s mergePeaks tool (mergePeaks {condition 1} {condition 2} -strand > output.txt) (Heinz et al. 2010). Raw read counts were then quantified for the each of the comparisons between conditions (annotatePeaks.pl {TSR} {genome} -gtf {antn} - strand + -fragLength 1 -raw -d {tag directories} > output.txt). The resulting output was then analyzed using DESeq2 to calculate the rlog variance stabilized counts and identify differentially regulated TSRs (Love et al. 2014).

### Read histograms

RefSeq were extracted from .gtf files using “parseGTF.pl {species gtf} tss > {species}_refSeq.tss.txt”. Histograms showing experimental TSS to RefSeq TSS were created using “annotatePeaks.pl refSeq [genome] -p experimentalTSS -size 500 -hist 1 -strand + > output.tsv” from HOMER (Heinz et al. 2010).

### Motif analysis

The analyis of the core promoter elements (the TATA box and the Initiator) for our experimental TSS and RefSeq were performed using HOMER’s annotatePeaks.pl tool (annotatePeaks.pl {TSS} [genome] -size 150 -hist 1 -m {motif} > output.txt) (Heinz et al. 2010). This tool was also used to find the nucleotide frequency plots (annotatePeaks.pl {TSS} [genome] -size 1000 -hist 1 -di > output.txt).

### Modified bison GTF

The first column of the bison gtf file underwent an ID update to achieve consistency with the fasta genome. A key was generated, linking the old IDs to their new counterparts in the genome (Supplemental Data X). Chromosome IDs (1-29, Y) were directly replaced with their corresponding accession numbers found in the original study’s NCBI BioProject repository (<u>PRJNA677946</u>) (Oppenheimer et al. 2021). For IDs labeled “scaffold_XXX,” a specific transformation was applied: 10,000,000.1 was added to the number following the underscore, and the resulting number was prefixed with “JAEQBK0.” The modified bison gtf (modified_GCA_018282365.1) was then created using the following custom code.

`name_key_df = pd.read_csv(“name_key.csv”, dtype={“old_id”: “string”})`

`name_key = dict(zip(name_key_df[“old_id”], name_key_df[“new_id”]))`

`cols = [’gtf_id’, ‘source’, ‘feature’, ‘start’, ‘end’, ‘score’, ‘strand’, ‘frame’, ‘attributes’]`

`gtf = pd.read_csv(“Bison_bison_liftoff.ARS-UCSC_bison1.0.gtf”, sep = ‘\t’, header = None, names = cols, dtype={“gtf_id”: “string”})`

`modified_gtf = gtf.assign(gtf_id=gtf[“gtf_id”].map(name_key)) modified_gtf.to_csv(’modified_Bison_bison_liftoff.ARS-UCSC_bison1.0.gtf’, sep=’\t’, index=False, header=False, quoting=3)`

## DATA DEPOSITION

All next-generation-sequencing (NGS) data are available at NCBI Gene Expression Omnibus with the accession number GSE267848. Other data that support the findings of this study are available from the corresponding author upon request.

## SUPPLEMENTAL MATERIAL

Supplemental material is available for this article.

## ACKNOWLEDGMENTS

We thank Dr. Tony M. Mertz and Steven A. Roberts for reagents, and the Molecular Biology and Genomics Core at Washington State University for Illumina NextSeq2000 sequencing. This manuscript includes data generated at the UC San Diego IGM Genomics Center utilizing an Illumina NovaSeq 6000 that was purchased with funding from a National Institutes of Health SIG grant (#S10 OD026929). Research was supported by the National Institutes of Health (NIH) grant R00 GM135515 and WSU startup funds to S.H.D., Agricultural Research Service (ARS), U.S. Department of Agriculture (USDA) grant number CWU 2090-32000-045-00D to C.W.C. supported animal research involving the cattle and bison from which blood samples were obtained. D.D.M was supported by the Oak Ridge Institute for Science and Education (ORISE) contract number DE-SC0014664.

## AUTHOR CONTRIBUTIONS

A.L.M, A.M.B, J.Z. and S.H.D. oversaw the overall design, conceptualization, and execution of the project. The experiments were performed by A.M.B. and M.I.S. The computational analyses were performed by A.L.M, I.M.B., and S.H.D. were primarily responsible for writing the manuscript. X.Q. provided sequencing methodology, D.D.W and C.W.C reagents and expertise. All authors revised and approved the final manuscript.

## ETHICAL APPROVAL

Blood collection from experimental animals followed protocols approved by the Institutional Animal Care and Use Committee of Washington State University (Animal Subject Approval Form #7080 for cattle) or University of Wyoming (Protocol # 20230201BW00287-01 for bison).

## SUPPLEMENTAL MATERIAL

**Supplemental Table S1**

Sequencing quality statistics across both sequencing platforms for all samples. Statistics were summarized using Bases2Fastq, FastQC (to assess %GC, duplicate rate, avg length, and median length), and Seqtk (to assess percent Q30, percent Q40, mean quality score).

## Supplemental Protocol

Detailed description of the AVITI short fragment sequencing procedure.

---

**Figure S1:**
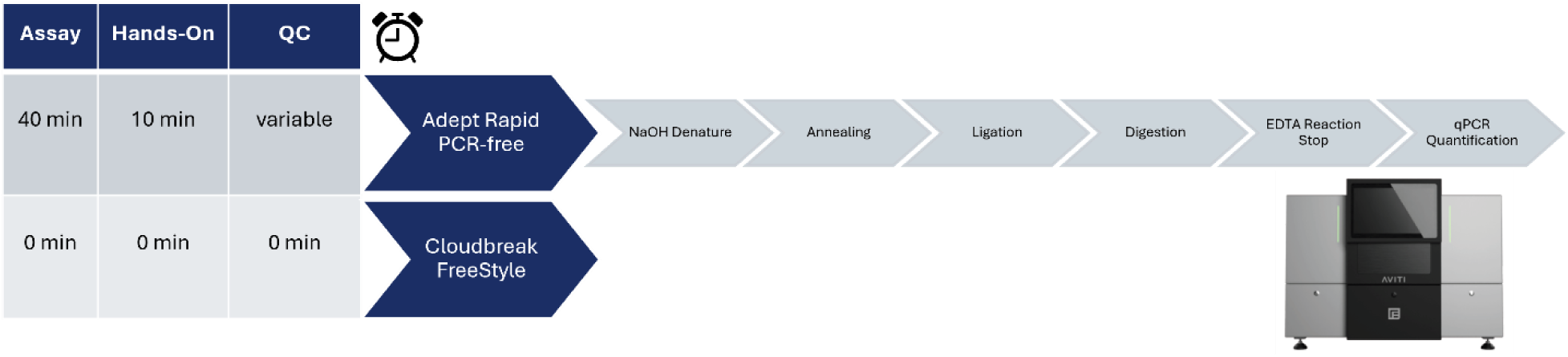
Overview of optimized small DNA fragment sequencing workflow using AVITI. Circularization methods for AVITI sequencing. All assays require quality standard sequencing library control prior to beginning. Adept Rapid PCR-free implements benchtop conversion strategies and additional QC prior to sequencing. Cloudbreak Freestyle does not require any additional processing before sequencing.

**Figure S2:**
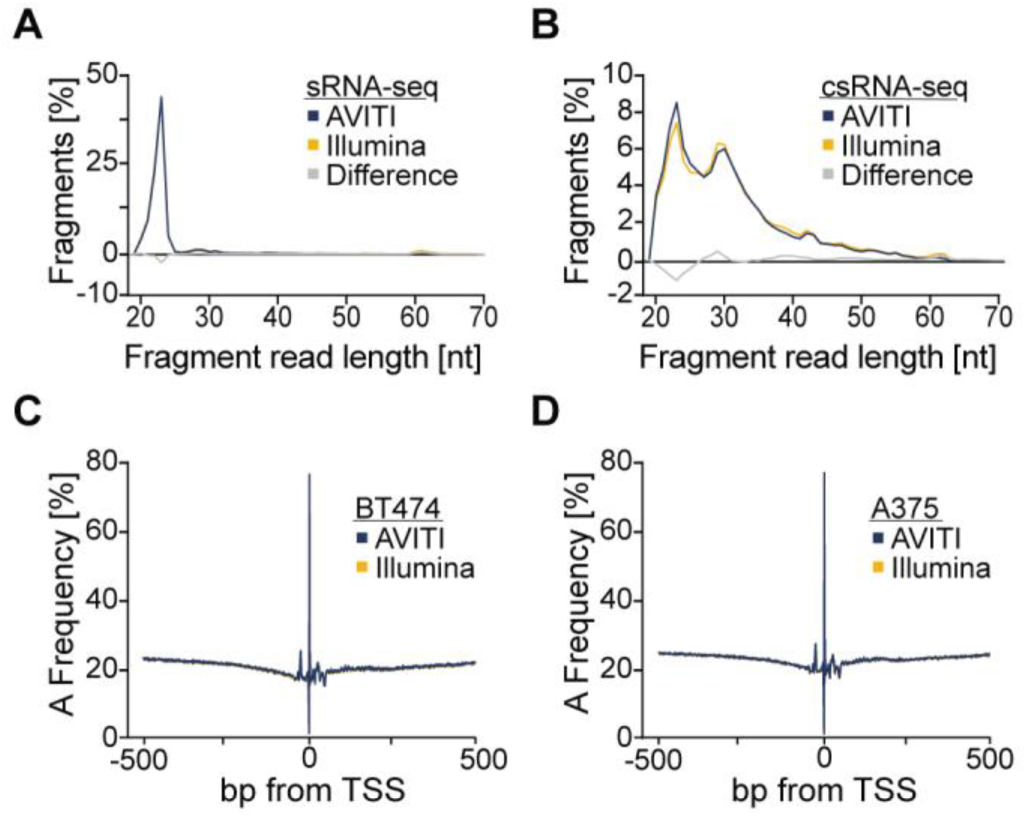
Additional comparisons of fragment length and nucleotide biases **A**. Read length distribution plots of BT474 small RNAs sequenced on the Illumina and AVITI platform using the Cloudbreak Freestyle method. The area under each line sums to a total of 100%, except the grey line which denotes the difference between Illumina and AVITI. **B**. Read length distribution plots of BT474 capped small RNAs. **C**. A nucleotide frequency plot of TSSs from Illumina and AVITI for BT474 and **D.** A375 human cancer cells.

**Figure S3:**
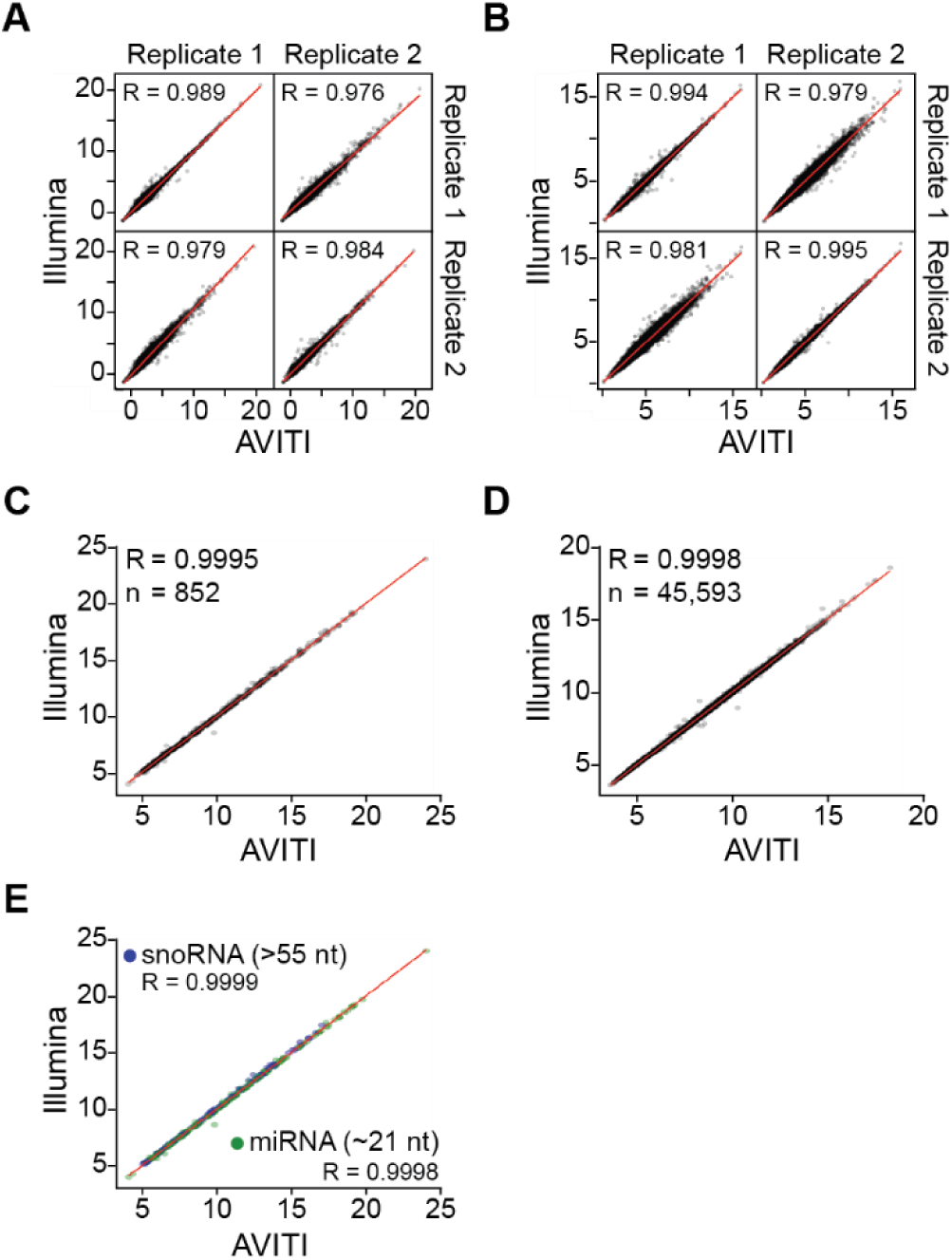
Correlation among libraries sequenced using AVITI and Illumina. **A.** Scatterplot comparing the expression level among replicates of A375 small RNAs and **B.** capped small RNAs using the Illumina and AVITI platform. **C.** Scatterplot comparing the expression level of BT474 small RNAs and **D**. capped small RNAs using the Illumina and AVITI platform. **E**. Comparison of the detection of BT474 small RNA types of different lengths (miRNAs: 21-24; snoRNAs: 55-61).

**Figure S4:**
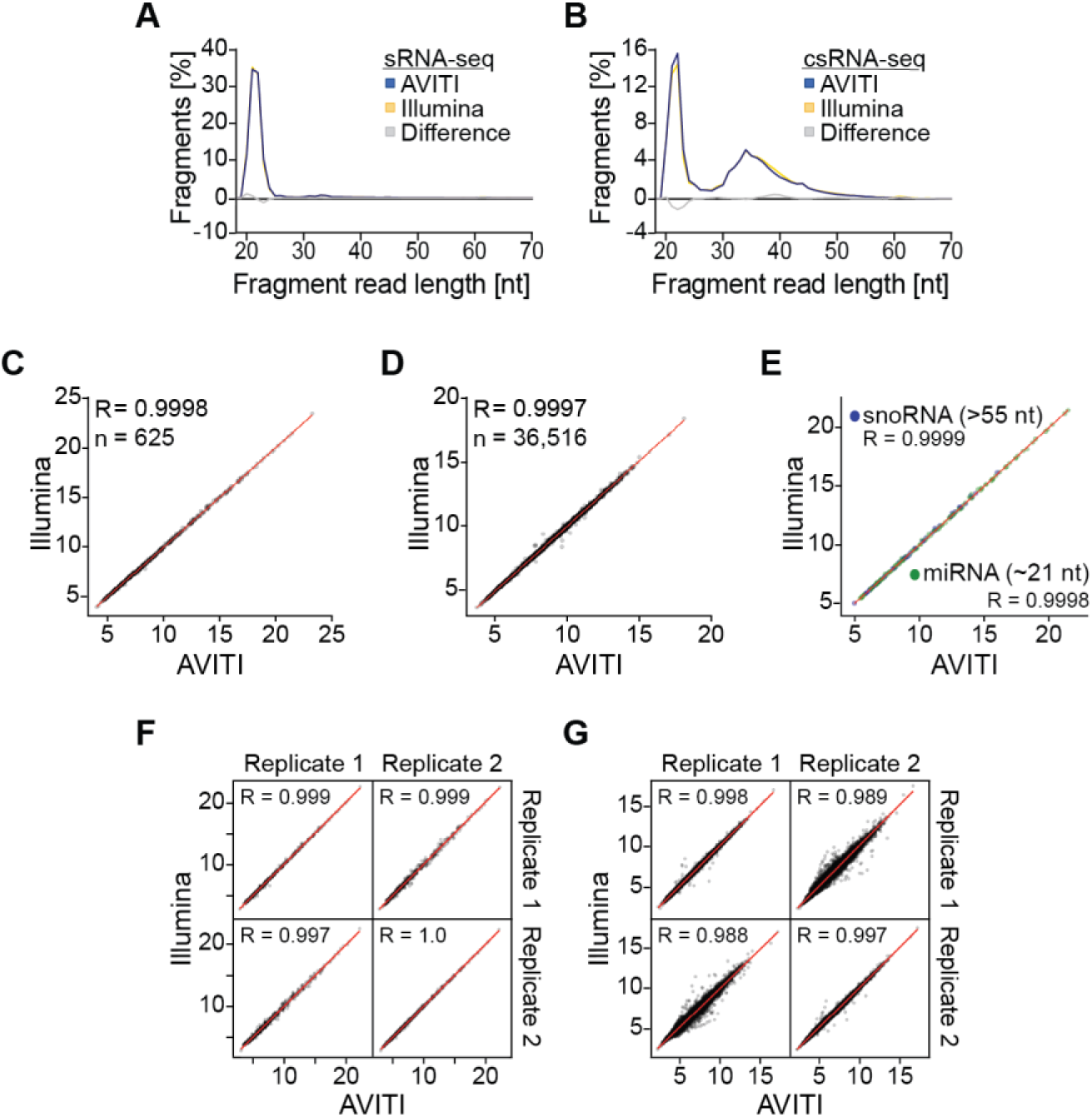
Correlation among libraries generated for cattle sequenced using AVITI and Illumina. **A.** Read length distribution plots of cattle small RNAs sequenced on the Illumina and AVITI platform. Each graph sums up to 100% except the grey graph which denotes the difference between Illumina and AVITI. **B.** Read length distribution plots of cattle capped small RNAs. **C**. Scatterplot comparing the expression level of small RNAs and **D**. capped small RNAs using the Illumina and AVITI platform. **E**. Comparison of the detection of small RNA types of different lengths (miRNAs: 21-24; snoRNAs: 55-61). **F.** Scatterplot comparing the expression level among replicates of cattle small RNAs and **G.** capped small RNAs using the Illumina and AVITI platform.

**Figure S5:**
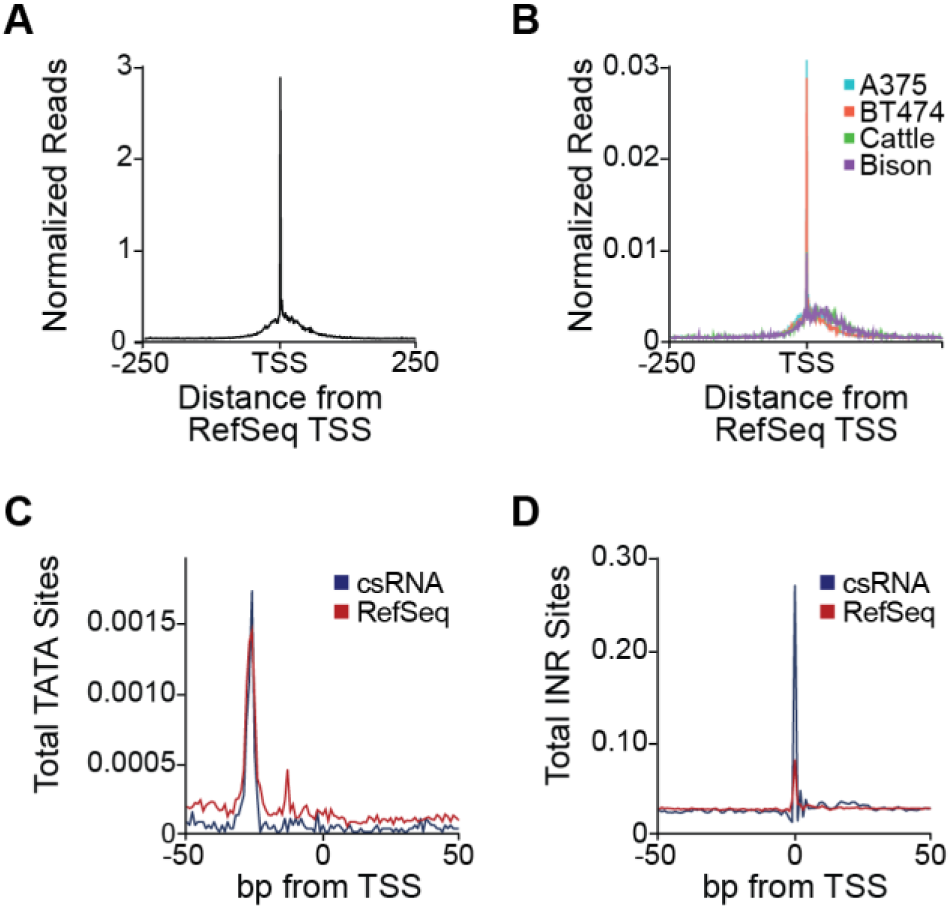
csRNA-seq could be used to improve 5’ annotations of livestock genes. **A.** Comparison of experimentally defined TSSs from BT474 by csRNA-seq relative to the Human RefSeq annotation and **B.** all comparison of experimentally defined TSS from the respective RefSeq. **C.** Comparison of the frequency of TATA box sites and **D.** Initiator sites per 1000 bp between our experimental TSS and RefSeq for BT474.

**Figure S6:**
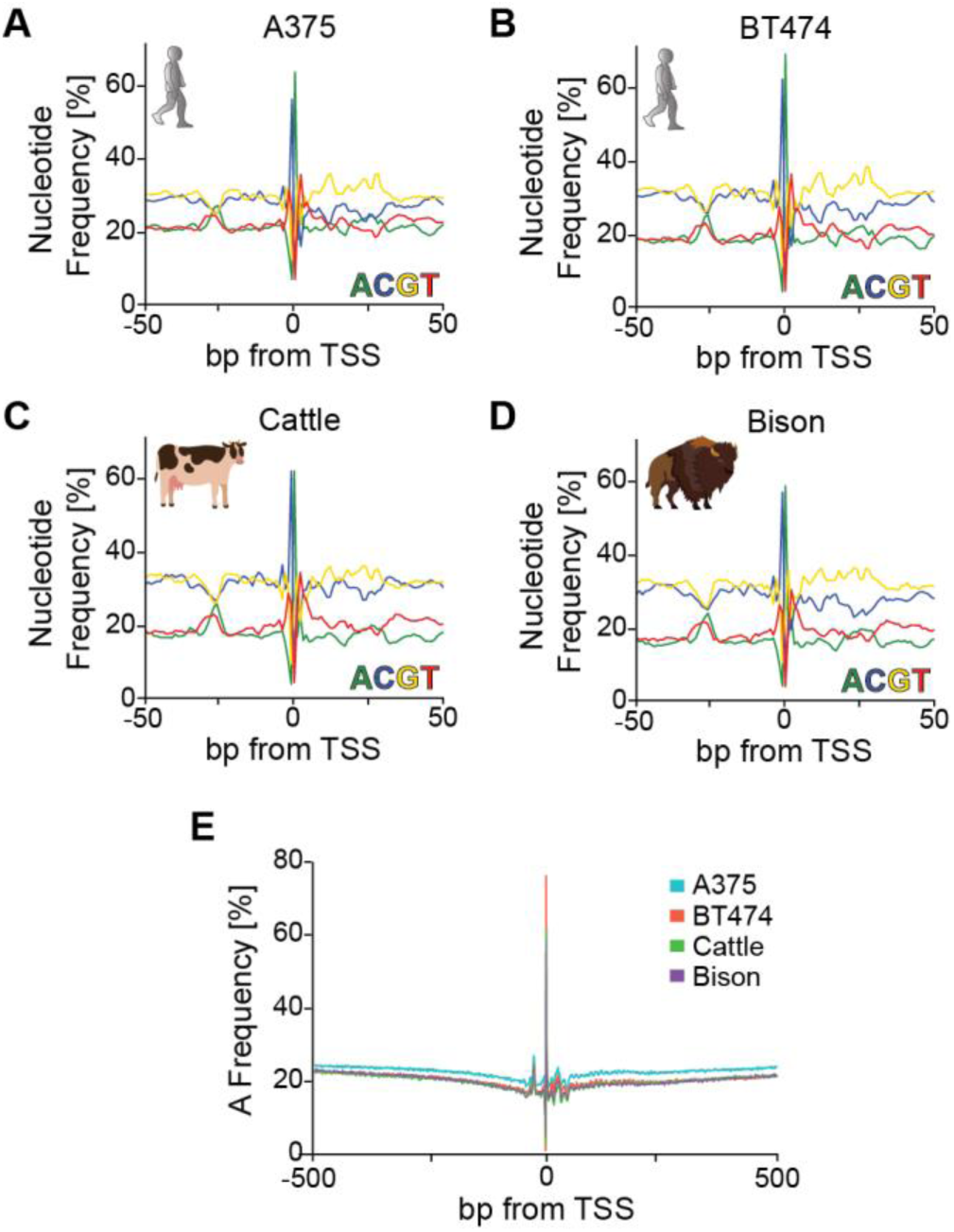
Nucleotide frequencies near human, cattle and bison transcription start sites experimentally defined by csRNA-seq. **A.** Nucleotide frequency plots of TSSs in human A375 cancer cells, **B.** human BT474 cancer cells, **C.** cattle, and **D.** bison. **E.** Combined A nucleotide frequency plot for each species and cell line.

## Element Adept^™^ Library Compatibility Workflow

### Introduction

The Element Adept Library Compatibility Workflow adapts linear libraries prepared with third-party kits for sequencing on the Element AVITI™ System. The Adept Rapid PCR-Free Protocol uses the Element Adept Library Compatibility Kit v1.1 to circularize up to 24 reactions. Each reaction supports input of one linear library or a pool of indexed linear libraries.

The protocol starts with denaturing a linear library into single strands. The library is then annealed to splint oligos, which introduce the Element surface primer binding sequences. A ligation reaction circularizes the library, followed by enzymatic digestion for cleanup.

**Figure 1a:**
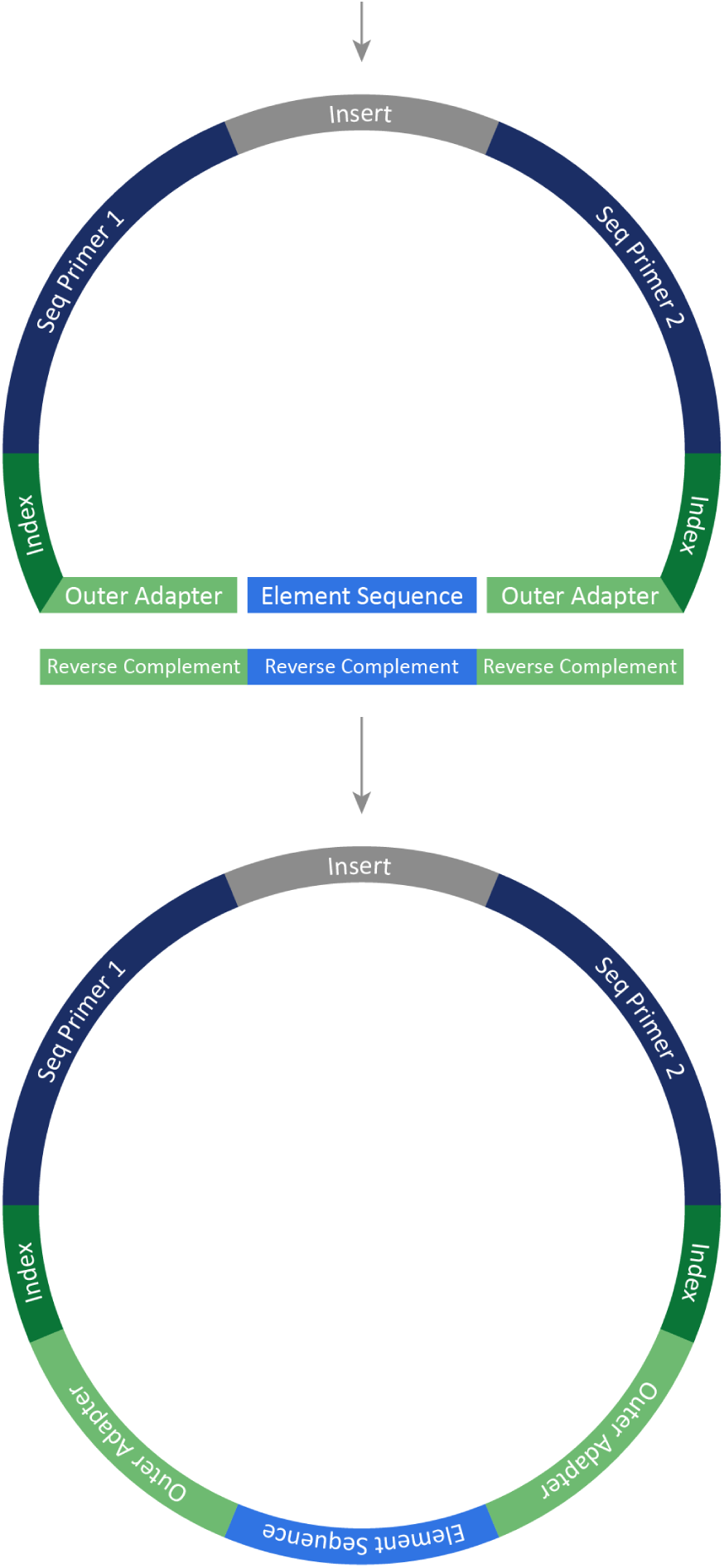
Circularization of a linear library.

#### Sequencing Compatibility

Adept libraries are *only* compatible with Cloudbreak™ sequencing kits on the Element AVITI System. If you are using Cloudbreak Freestyle™ sequencing kits, converting your library with the Adept Workflow is not required. For more compatibility information, see [go.elembio.link/product-compatibility](https://www.elementbiosciences.com/product-compatibility).

#### Supported Protocols

The Adept Workflow supports multiple kits and protocols. The compatibility kit generates a circular library and the PCR-plus kit generates a linear library that is circularized onboard the instrument. Thus, the AVITI System sequences a circular library regardless of whether you load a linear or circular library.

This guide documents the Adept Rapid PCR-Free Protocol. To follow a different protocol, reference the applicable user guide.

| Protocol | Kit | User Guide |
| --- | --- | --- |
| Adept Standard | Element Adept Library Compatibility Kit v1.1 | *Element Adept Library Compatibility Workflow User Guide for the Standard Protocol (MA-00001)* |
| Adept Rapid PCR-Free | Element Adept Library Compatibility Kit v1.1 | Current user guide |
| Adept Rapid PCR-Plus | Adept Rapid PCR-Plus Kit | *Element Adept Library Compatibility Workflow User Guide for the PCR-Plus Protocol (MA-00040)* |

#### Library Compatibility

The Adept Rapid PCR-Free Protocol supports libraries prepared with the library prep and index kits listed at <u>go.elembio.link/compatible</u>. If your kit is not supported, follow the Adept PCR-Plus Protocol.

The following factors make a library unsupported:

- Truncated ends
- End-blocked libraries
- Additional bases at the end, such as a polymerase-generated adenine (A) overhang

PCR-free libraries constructed with IDT xGen UDI-UMI Adapters are not compatible with the Adept Rapid PCR-Free Protocol. IDT xGen UDI-UMI Adapters have the insert flipped to the opposite strand as compared to standard libraries. For libraries prepared with these adapters, follow the Adept PCR-Plus Protocol. For more compatibility information, see <u>go.elembio.link/product-compatibility</u>.

#### Quantification Method

Quantifying the final library ensures the appropriate input for sequencing. By detecting only circular library, quantitative PCR (qPCR) ensures consistent and accurate quantification of libraries prepared with the Element Adept Library Compatibility Kit v1.1.

Qubit is an alternative quantification method that requires testing and modifications. For more information, see the *Accurate Quantification of Circular Libraries for Sequencing on the Element AVITI System Technical Note (LT-00009)*.

#### Low-Diversity Amplicon Library

When preparing a low-diversity amplicon library, such as 16S, for sequencing with a 2 x 300 kit, meet the following requirements:

- An insert size of > 200 bp
- High plexity of ≥ 64 unique dual indexed (UDI) libraries
- A 1–5% spike-in of PhiX Control Library

#### Custom Primers

The AVITI System accepts custom primers for any Adept library. However, custom primers require special consideration and planning. To make sure a run with custom primers meets specifications, contact Element Technical Support early in experiment planning.

Technical support can also help determine whether your library requires custom primers.

#### Workflow Summary

The following figure summarizes the protocol, which takes 40 minutes, including 10 minutes of hands-on time. All durations are approximate and depend on lab-specific factors.

**Figure 2:**
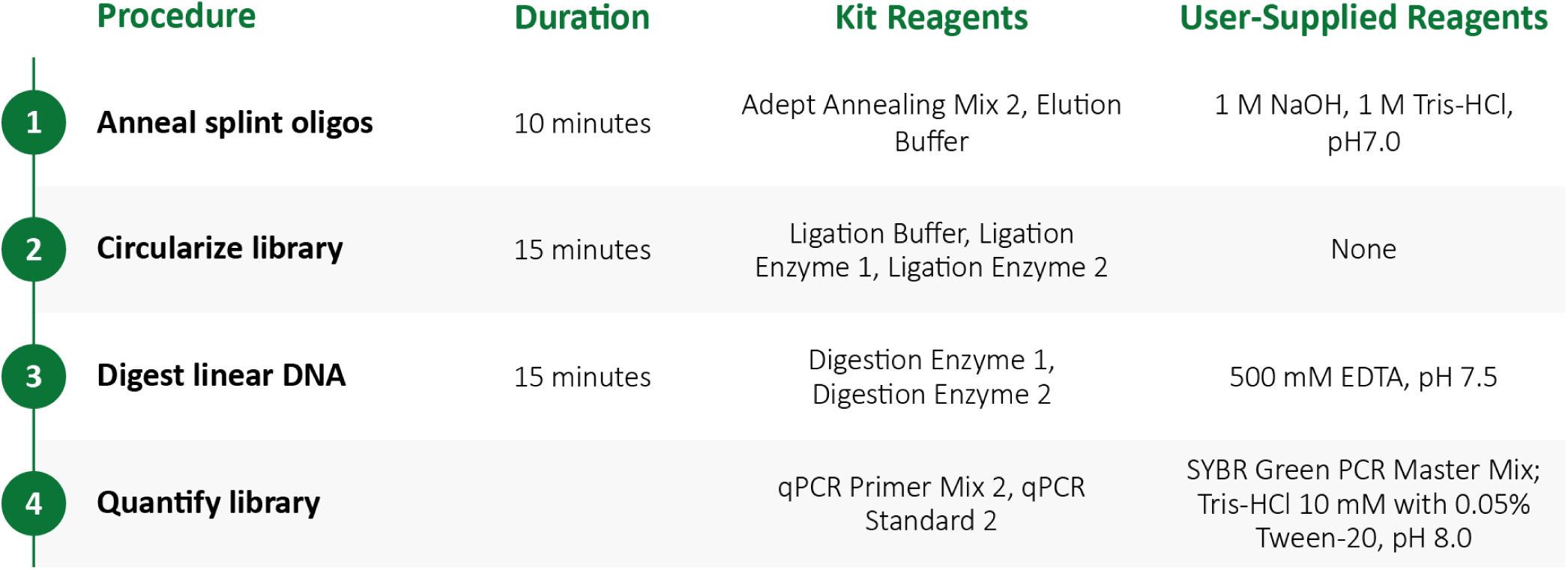
Adept Rapid PCR-Free Protocol

#### Safety Data Sheets

When using the Element Adept Library Compatibility Kit v1.1 and other reagents, always wear personal protective equipment (PPE): a lab coat, powder-free disposable gloves, and protective goggles. Review the safety data sheets (SDS) for chemical properties. The SDS inform safety, disposal, and hazards for your region and are available at <u>elementbiosciences.com/resources</u>.

### Kit Contents and Storage

The Element Adept Library Compatibility Kit v1.1 is packaged in one box and shipped on dry ice. When you receive your kit, promptly store reagents at the proper temperature. Reference reagent labels for fill volumes.

In addition to the kit, the protocol requires the user-supplied materials listed in the following sections. The protocol specifies processing libraries in a plate, but you can substitute tubes.

| Reagent | Quantity | Cap Color | Storage Temperature |
| --- | --- | --- | --- |
| Adept Annealing Mix 2 | 1 | Green | -25°C to -15°C |
| Digestion Enzyme 1 | 1 | Clear | -25°C to -15°C |
| Digestion Enzyme 2 | 1 | Clear | -25°C to -15°C |
| Elution Buffer | 1 | Clear | -25°C to -15°C |
| Ligation Buffer | 2 | Clear | -25°C to -15°C |
| Ligation Enzyme 1 | 1 | Clear | -25°C to -15°C |
| Ligation Enzyme 2 | 1 | Clear | -25°C to -15°C |
| qPCR Primer Mix 2 | 1 | Clear | -25°C to -15°C |
| qPCR Standard 2 | 1 | Clear | -25°C to -15°C |

#### User-Supplied Consumables

| Supplier | Consumable | Catalog # |
| --- | --- | --- |
| General lab supplier | 96-well PCR plates | Not applicable |
|  | 96-well qPCR plates | Not applicable |
|  | Absolute ethanol | Not applicable |
|  | Filtered pipette tips | Not applicable |
|  | Nuclease-free water | Not applicable |
|  | Sample purification beads | Not applicable<sup>1</sup> |
| Agilent | High Sensitivity D5000 Reagents | Part # 5067-5593<sup>2</sup> |
|  | High Sensitivity D5000 ScreenTape | Part # 5067-5592<sup>2</sup> |
| Bio-Rad | Microseal 'B' Film, adhesive | Catalog # MSB1001<sup>2</sup> |
|  | Microseal 'C' Film, optical | Catalog # MSC1001<sup>2</sup> |
| Eppendorf | DNA LoBind Tubes, 1.5 ml | Catalog # 022431021 |
| Teknova | 1 M Tris-HCl, pH 7.0 | SKU # T1070<sup>2</sup> |
|  | 1 N Sodium Hydroxide (NaOH) | SKU # H0224<sup>2</sup> |
|  | 10 mM Tris-HCl with 0.05% Tween-20, pH 8.0 | SKU # T1485<sup>2</sup> |
|  | 500 mM EDTA, pH 7.5 | SKU # E0375<sup>2</sup> |
| Thermo Fisher Scientific | 96-well 0.8 ml deepwell storage plates | Catalog # AB0765<sup>2,3</sup> |
|  | Either kit:<br>• Qubit dsDNA BR Assay Kit<br>• Qubit dsDNA HS Assay Kit | The corresponding catalog #:<br>• Q33216<br>• Q33238 |
|  | SYBR Green PCR Master Mix | Catalog # 4364346<sup>2</sup> |

<sup>1</sup> Element has validated SPRIselect, 60 ml (Beckman Coulter, catalog # B23318).
<sup>2</sup> Consumables that you have tested and demonstrate equivalent performance are acceptable.
<sup>3</sup> Deepwell plates facilitate cleanup procedures. When cleaning up libraries, you can instead use 0.2 ml tubes or strip tubes.

#### User-Supplied Equipment

| Supplier | Equipment | Catalog #<sup>1</sup> |
| --- | --- | --- |
| General lab supplier | Centrifuge, multipurpose | Not applicable |
|  | Ice bucket | Not applicable |
|  | Pipettes, single- or multi-channel | Not applicable |
|  | Vortex mixer | Not applicable |
|  | [Optional] Speed vac | Not applicable |
| Agilent | Either system:<br>• 4150 TapeStation System<br>• 4200 TapeStation System | The corresponding part #:<br>• G2992AA<br>• G2991BA |
| Bio-Rad | CFX96 Touch Real-Time PCR Detection System | Catalog # 1845096<sup>1</sup> |
|  | Either thermal cycler:<br>• C1000 Touch Thermal Cycler<br>• T100 Thermal Cycler | The corresponding catalog #:<br>• Catalog # 1851197<br>• Catalog # 1861096 |
| Thermo Fisher Scientific | The applicable magnet:<br>• DynaMag-96 Side Magnet for cleanup with tubes<br>• Magnetic Stand-96 for cleanup with plates | The corresponding catalog #:<br>• 12331D<br>• AM10027 |
|  | Either fluorometer:<br>• Qubit 3 Fluorometer<br>• Qubit 4 Fluorometer | The corresponding catalog #:<br>• Q33216<br>• Q33238 |
| QInstruments | [Optional] BioShake XP | Order # 1808-0505 |

<sup>1</sup> Any equipment that you have tested and demonstrates equivalent performance is acceptable.

### Input Requirements

The Adept Workflow supports a double-stranded DNA (dsDNA) linear library prepared per third-party instructions. Accordingly, you must prepare a linear library and perform a quality control (QC) check *before* starting end polishing or any of the protocols.

Prepare the linear library from RNA, complementary DNA (cDNA), or genomic DNA (gDNA). For more information, see <u>Library Compatibility on page 5</u>.

#### Library Amount

The Element Adept Library Compatibility Kit v1.1 accepts input of 4.2–20.8 nM in 24 µl low TE buffer or similar solution, which is equivalent to 0.1–0.5 pmol. The concentration of ethylenediaminetetraacetic acid (EDTA) in the library cannot exceed 1 mM.

Determine the input library concentration with a Qubit fluorometer, qPCR, or equivalent, following supplier instructions. When reducing the input amount to < 0.5 pmol, accurate quantification is crucial.

#### Fragment Size

Use a 4150 TapeStation System, 4200 TapeStation System, or equivalent instrument to qualify the input library and identify short byproducts in electropherograms. Set the region to > 175 bp and determine the average fragment size. Library portions that contain > 1000 bp sequences might impact Q30 scores and require adjustment for density.

Avoid significant amounts of adapter dimer or other short byproducts (< 175 bp). If the input library contains short byproducts, Element recommends additional cleanup using sample purification beads. Reassess the purified library with the TapeStation System to confirm byproduct removal, and then requantify.

**Figure 3:**
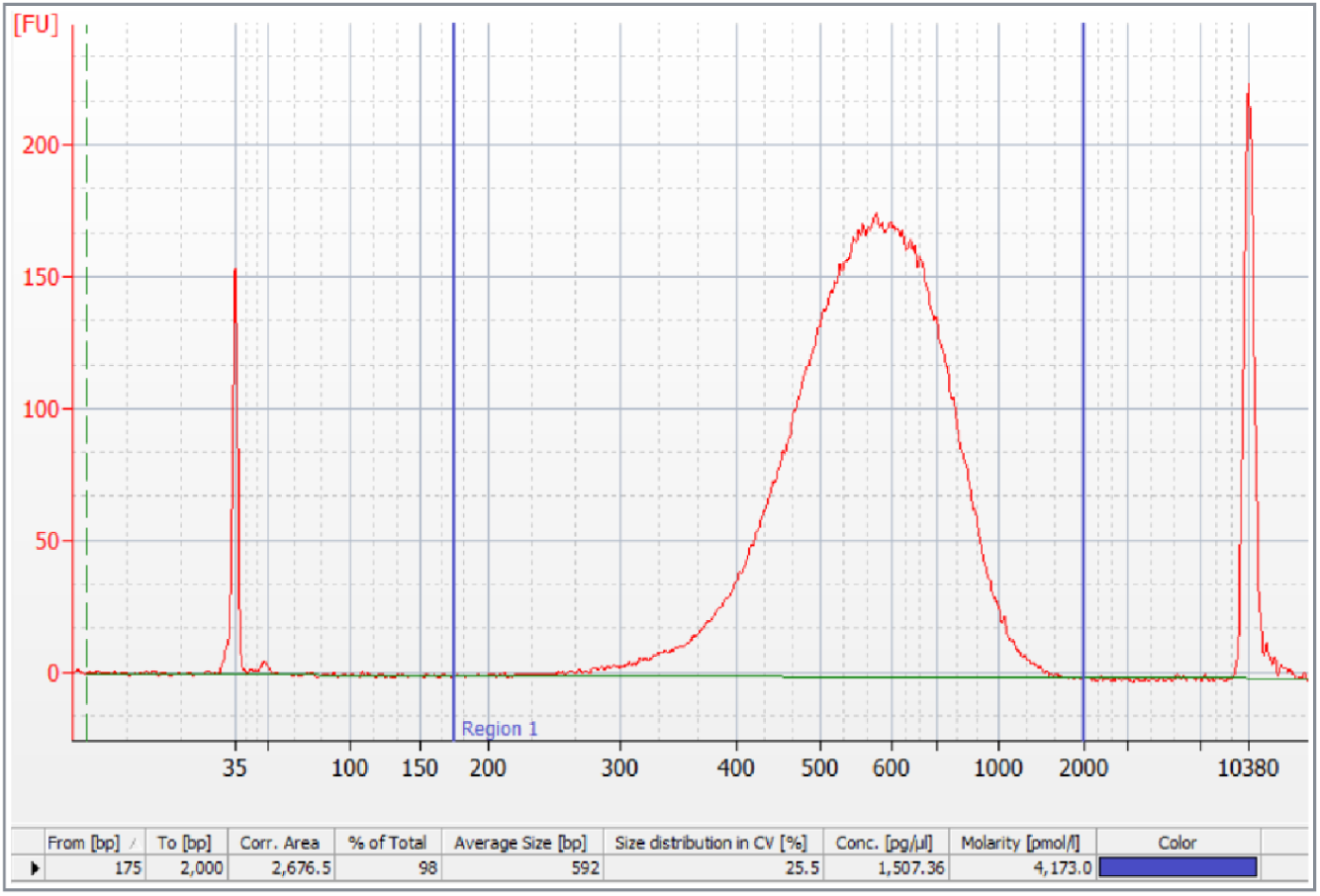
Example Bioanalyzer trace showing an average fragment size of 592 bp

#### Pooling Guidelines

An Adept reaction processes one linear library or a pool of indexed linear libraries. When pooling, uniquely index each library in the pool and apply the following criteria to pool libraries with similar characteristics:

- Pool libraries that require the same run parameters.
- Do not pool Adept libraries with Elevate™ libraries.
- Balance the concentrations of libraries in a pool based on the throughput requirements for each sample. To maintain balance after library prep, make sure the libraries have similar size distributions.
- Review the *Element AVITI System User Guide (MA-00008)* for guidance on using the PhiX Control Library, which can improve color and nucleotide balancing and library complexity. Certain experiments require a spike-in.

### Adept Rapid PCR-Free Protocol

Follow the protocol steps in the order listed using specified volumes and durations. Proceed immediately from one step to the next.

To avoid cross-contamination, use filtered pipette tips throughout the protocol. When adding or transferring reagents and libraries, change pipette tips between each reagent and each library.

#### Prepare Reagents

Reagent preparation is a preliminary procedure. Start 20–30 minutes before proceeding with the protocol.

1. Make sure that you have all Element- and user-supplied consumables. For lists, see *<u>Kit Contents and Storage</u>* <u>on page 7</u>.
2. Remove the Element Adept Library Compatibility Kit v1.1 reagents from -25°C to -15°C storage.

» Avoid unnecessary freeze-thaw cycles of Ligation Buffer:

◦ If you are preparing ≤ 12 reactions, remove one tube.
◦ If you are preparing 13–24 reactions, remove two tubes.
» If you are quantifying libraries immediately after circularization, remove qPCR Standard 2 and qPCR Primer Mix 2.
3. If necessary, remove the library from -25°C to -15°C storage and thaw on ice.
4. Fully thaw the reagents on ice. Keep on ice.

#### Anneal Splint Oligos

The anneal splint oligo procedure denatures the input linear library with NaOH and anneals splint oligos.

1. Gather the following consumables:

» 1.5 ml DNA LoBind tube
» 96-well PCR plate
» Microseal ‘B’
» 1 M NaOH
» 1 M Tris-HCl, pH 7.0
» Adept Annealing Mix 2 (green cap)
» Elution Buffer
» Linear library
2. Make sure a dsDNA linear library is prepared per supplier instructions. Prepare 24 µl 4.2–20.8 nM linear library or indexed linear library pool so the total input amount is ∼0.1–0.5 pmol.

» If the volume is < 24 µl, add Elution Buffer to reach 24 µl.
» If the volume is > 24 µl, use a speed vac or validated bead-based method to concentrate the library to 24 µl.
3. Make sure that Adept Annealing Mix 2 is fully thawed.
4. Vortex Adept Annealing Mix 2 to mix and briefly centrifuge.
5. Add 24 µl library to each well of a new PCR plate.
6. Add 3 µl 1 M NaOH to each well.
7. Mix using either method:

» Seal the plate and vortex.
» Set a pipette to 19 µl, pipette each reaction 10 times to mix, and seal the plate.
8. Incubate at room temperature for 5 minutes.
9. During incubation, combine the following reagents to prepare fresh master mix, allowing 10–15% overage. Set a pipette to 70% of the master mix volume and pipette 10 times on ice to mix.
10. After incubation, add 16 µl master mix to each well.
11. Mix using either method:

» Seal the plate and vortex.
» Set a pipette to 30 µl, pipette each reaction 10 times to mix, and seal the plate.
12. Briefly centrifuge the plate.
13. Place the plate in the thermal cycler.
14. Run the following ∼5-minute program:
15. Remove the plate from the thermal cycler.
16. Briefly centrifuge the plate and immediately proceed.

| Reagent | Volume per Reaction (μl) |
| --- | --- |
| 1 M Tris-HCl, pH 7.0 | 3 |
| Adept Annealing Mix 2 | 13 |
| Total | 16 |

| Step | Temperature | Time |
| --- | --- | --- |
| Volume set to 43 μl | | |
| Lid set to 45°C |  |  |
| 1 | 37°C | 5 minutes |
| 2 | 37°C | Hold |

#### Circularize Library

The circularize library procedure phosphorylates the 5′ end of the linear library and uses a ligation reaction to produce a circular library.

1. Gather the following consumables:

» 1.5 ml DNA LoBind tube
» Microseal ‘B’
» Ligation Buffer
» Ligation Enzyme 1
» Ligation Enzyme 2
2. Make sure that all reagents are fully thawed.
3. Gently flick Ligation Enzyme 1 and Ligation Enzyme 2 to mix and briefly centrifuge. Place on ice.
4. Vortex Ligation Buffer to mix and briefly centrifuge.
5. Combine the following reagents to prepare fresh master mix, allowing 10–15% overage. Set a pipette to 70% of the master mix volume and pipette 10 times on ice to mix.
6. Add 7 µl master mix to each reaction.
7. Set a pipette to 38 µl and pipette each reaction 10 times to mix.
8. Seal the plate and briefly centrifuge.
9. Place the plate in the thermal cycler.
10. Run the following ∼10-minute program.
11. Remove the plate from the thermal cycler.
12. Briefly centrifuge the plate and immediately proceed.

| Reagent | Volume per Reaction (μl) |
| --- | --- |
| Ligation Buffer | 5 |
| Ligation Enzyme 1 | 1 |
| Ligation Enzyme 2 | 1 |
| Total | 7 |

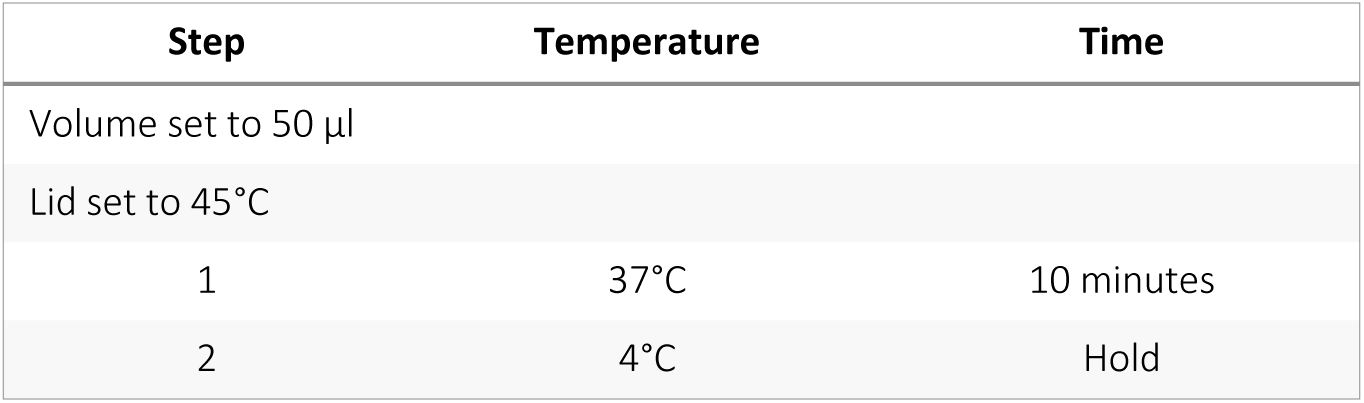

#### Digest Linear DNA

The digestion procedure removes carryover linear DNA.

1. Gather the following consumables:

» DNA LoBind tubes
» Microseal ‘B’
» 500 mM EDTA, pH 7.5 (EDTA)
» Digestion Enzyme 1
» Digestion Enzyme 2
2. Gently flick Digestion Enzyme 1 and Digestion Enzyme 2 and briefly centrifuge. Place on ice.
3. Combine the following reagents to prepare fresh master mix, allowing 10–15% overage. Set a pipette to 70% of the master mix volume and pipette 10 times on ice to mix.
4. Add 4 µl master mix to each reaction.
5. Set a pipette to 38 µl and pipette each reaction 10 times to mix.
6. Seal the plate and briefly centrifuge.
7. Place the plate in the thermal cycler.
8. Run the following ∼10-minute program:
9. Remove the plate from the thermal cycler.
10. Vortex EDTA to mix and briefly centrifuge.
11. Add 2 µl EDTA to each reaction to neutralize.
12. Set a pipette to 39 µl and pipette each reaction 10 times to mix.
13. Seal the plate and briefly centrifuge.
14. Transfer each reaction (56 µl) to a new DNA LoBind tube.

—The tubes contain the final circular libraries.—
15. If you are not immediately quantifying or sequencing, cap the tubes and store at -25°C to -15°C for ≤ 15 days.

—The *Element* AVITI *System User Guide (MA-00008)* contains sequencing instructions, including diluting the library to the loading concentration.—

| Component | Volume per Reaction (μl) |
| --- | --- |
| Digestion Enzyme 1 | 2 |
| Digestion Enzyme 2 | 2 |
| Total | 4 |

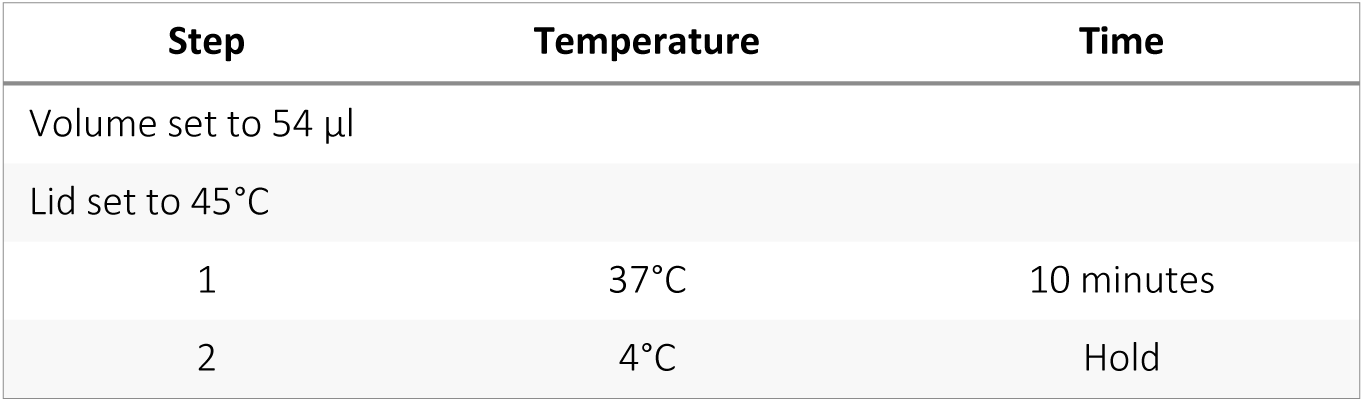

| Step | Temperature | Time |
| --- | --- | --- |
| Volume set to 54 μl |  |  |
| Lid set to 45°C |  |  |
| 1 | 37°C | 10 minutes |
| 2 | 4°C | Hold |

#### Quantify Library

The quantify library procedure uses qPCR to generate PCR amplicons over the ligated junctions and quantify a portion of the library in preparation for sequencing. The procedure requires standard and library dilutions run in triplicate qPCR reactions.

Each qPCR reaction is 10 µl and includes the following components:

- 1 µl 10x qPCR Primer Mix 2
- 4 µl standard, library, or any positive or negative control diluted to assay-appropriate levels
- 5 µl 2x SYBR Green PCR Master Mix

#### Prepare Dilutions

1. Gather the following consumables:

» 1.5 ml DNA LoBind tube
» 96-well qPCR-compatible plate (assay plate)
» Microseal ‘C’
» 10 mM Tris-HCl with 0.05% Tween-20, pH 8.0 (dilution buffer)
» Circular library
» qPCR Primer Mix 2
» qPCR Standard 2
» SYBR Green PCR Master Mix
2. Prepare the library, qPCR Primer Mix 2, and qPCR Standard 2:

a. Thaw the library and reagents on ice.
b. Make sure the library and reagents are fully thawed.
c. Pulse vortex the library and reagents and briefly centrifuge.
3. Set aside ∼20 µl dilution buffer as a no-template control (NTC).
4. In a 1.5 ml DNA LoBind tube, combine the following reagents to prepare qPCR Standard 2.
5. Vortex the tube to mix and briefly centrifuge.
6. Label the tube **200 pM qPCR Standard 2**.
7. From the 200 pM qPCR Standard 2, make 1:10 serial dilutions to prepare the following standard dilutions.

—Each standard requires 12 µl for triplicate reactions.—
8. [Optional] Store unused 200 pM qPCR Standard 2 at -25°C to -15°C for ≤ 15 days. Avoid frequent freeze-thaw cycles.
9. Using two 1:100 dilutions, dilute 2 µl library 1:10,000 in dilution buffer. If your expected yield is lower or higher than the typical yield, adjust the dilution.

—Libraries diluted to ∼0.1–1 pM typically appear in the middle of the standard curve and provide the most accurate quantification. Proper dilution for library is 1:10,000.—
10. Return the remaining library to -25°C to -15°C storage.

| Reagent | Volume per Reaction (µl) |
| --- | --- |
| Dilution buffer | 18 |
| 2 nM qPCR Standard 2 | 2 |
| Total | 20 |

| Standard | Concentration (pM) |
| --- | --- |
| Std 1 | 20 |
| Std 2 | 2 |
| Std 3 | 0.2 |
| Std 4 | 0.02 |
| Std 5 | 0.002 |
| Std 6 | 0.0002 |

#### Prepare Master Mix and Assay Plate

1. Combine the following reagents to prepare fresh qPCR master mix with primers, allowing 10–15% overage.

» Set a pipette to 70% of the master mix volume and pipette the master mix 10 times on ice to mix.
» Prepare sufficient volume to run triplicate reactions of each NTC, standard dilution, and library dilution.
2. Add 6 µl qPCR master mix with primers to the desired wells of a new assay plate.
3. Add 4 µl NTC, standard dilutions, or library dilutions to wells containing qPCR master mix with primers.

—The assay volume is 10 µl per well. Mixing is not necessary.—
4. Repeat steps 2–3 to prepare triplicate reactions of each NTC, standard dilution, and library dilution.
5. Seal the plate and briefly centrifuge.

| Reagent | Starting Concentration | Volume per Reaction (µl) |
| --- | --- | --- |
| SYBR Green PCR Master Mix | 2x | 5 |
| qPCR Primer Mix 2 | 10x | 1 |
| Total |  | 6 |

#### Perform a qPCR Run

1. On the run setup page of the qPCR run software, edit the plate file:

a. Assign standard wells and set the corresponding concentrations.
b. Assign NTC wells.
c. Assign library wells and note the dilution factor.
d. Assign any reference libraries, positive controls, or negative controls to the appropriate wells.
2. Place the plate in the qPCR instrument.
3. Run the following > 1-hour program on the qPCR instrument. If you are not using the qPCR master mix and instrument specified in *<u>Kit Contents and Storage</u>* <u>on page 7</u>, adjust the program settings.
4. Follow vendor instructions to QC the run.

| Step | Setting |
| --- | --- |
| Volume set to 10 µl |  |
| Lid set to 105°C |  |
| Activation | 10 minutes at 95°C |
| PCR 40 cycles | 15 seconds at 95°C |
|  | 1 minute at 60°C |
|  | Plate read |
| Melt curve | 55°C to 95°C with increments of 1°C every 5 seconds |
|  | Plate read after each temperature step |

#### Analyze Results

1. Analyze the results of the qPCR run:

» Exclude the data described in *<u>Exclusion Criteria</u>*.
» Generate the standard curve as described in *<u>Standard Curve Criteria</u>*.
2. Determine the library dilution concentrations in pM using either method:

» Use the starting quantity (SQ) mean values reported by the qPCR instrument software.
» Calculate mean values based on the standard curve.
3. Calculate the initial library concentration based on dilutions and measured concentrations:

*input* library *concentration in nM* = (*fold dilution* * *quantification mean in pM*)/1000
—Size adjustment in quantification is not necessary.—

##### Exclusion Criteria

- Outliers on the amplification and melt curves and failed wells per third-party qPCR instructions.
- Outliers with a difference > 0.5 Cq for standard dilution, library dilution, and control wells running replicate reactions.
- Standard dilutions that amplify < 3 Cq values ahead of the NTC. Any exclusion except Std 6 (0.0002 pM) requires a rerun.
- Any libraries that had all dilutions amplify outside the standard range require a rerun with, if necessary, an adjusted fold of dilution. The standard curve, which is generated from the standard dilutions that passed the first two exclusion criteria, determines the dynamic range.

##### Standard Curve Criteria

Generate the standard curve from standard dilutions that passed the first two exclusion criteria and plot Cq values against the log concentration. When assessing the standard curve, apply the following passing criteria: standard dilution amplification is 90–110%, which is equivalent to a slope of -3.6 to -3.1, and R² > 0.99.

- If the amplification efficiency and R² value are out of range, reassess data points in the standard curve and exclude outliers. The remaining standard dilutions must have ≥ 3 dilution points. A dilution point is a set of duplicates or triplicates in one of the six standard dilutions.
- If the remaining standard dilutions do not have ≥ 3 dilution points, troubleshoot and repeat the qPCR run with freshly prepared dilutions and reagents. The resulting standard curve must meet all passing criteria.

### Technical Support

Visit the <u>User Documentation page</u> on the Element Biosciences website for additional guides and the most recent version of this guide. For technical assistance, contact Element Technical Support.

**Website:** www.elementbiosciences.com

**Telephone:** +1 866.ELEMBIO (+1 866.353.6246)

### Document History

| Document # | Date | Description of Change |
| --- | --- | --- |
| Document # MA-00033 Rev. E | March 2024 | • Added information about sequencing kit compatibility.<br>• Added requirements for low-diversity amplicon libraries. |
| Document # MA-00033 Rev. D | January 2024 | • Corrected footer. |
| Document # MA-00033 Rev. C | November 2023 | • Corrected missing volumes and ligation master mix volume table in Circularize Library.<br>• Corrected missing pipette setting and master mix reagent table in Prepare Master Mix and Assay Plate. |
| Document # MA-00033 Rev. B | November 2023 | • Corrected missing dilution values in Quantify Library, Prepare Dilutions. |
| Document # MA-00033 Rev. A | November 2023 | • Initial release |

